# CDK phospho-degron tags for dynamic protein regulation

**DOI:** 10.64898/2026.11.13.766399

**Authors:** Mihkel Örd, Nastassia Shtaida, Marko Lõoke, Alissa Agerova, Luka Bulatovic, Stepan Strelchenko, Ervin Valk, Ilona Faustova

## Abstract

Cyclin-dependent kinase (CDK) activity underpins cell cycle progression through dynamic and temporally ordered phosphorylation events. While transcriptional regulation has been extensively exploited in synthetic biology, the rational engineering of post-translational signaling circuits remains underdeveloped. Here, we introduce modular CDK-responsive phospho-degron tags derived from the Far1 regulatory region and demonstrate their ability to confer tunable, cell cycle-dependent control over protein stability in budding yeast. By systematically modifying cyclin docking motifs and Cks1 priming sites, we generate degron variants with distinct degradation kinetics and phosphorylation thresholds. These modules enable predictable modulation of transcription factor abundance and downstream gene expression. We further combine CDK-responsive degrons with phosphorylation-controlled localization modules for multi-layered regulation of sequential nuclear import and degradation. We encode multiple kinase activity thresholds within a single protein, demonstrating how complex phosphorylation logic can be compactly implemented. Finally, we show that degron performance is strongly influenced by culture conditions, particularly cell density and associated changes in CDK activity. Together, our results establish CDK-driven phospho-degrons as versatile building blocks for engineering dynamic protein regulation and provide a foundation for constructing synthetic post-translational signaling circuits.

## Introduction

Precise control over protein abundance and activity is a central objective in synthetic biology. Although transcriptional regulation has been widely used to construct synthetic gene circuits (Ellis et al. 2009; Khalil et al. 2012; Gander et al. 2017; Y. Chen et al. 2020; Elowitz and Leibler 2000), post-translational regulation provides complementary means of controlling protein levels with temporal precision (Grilly et al. 2007; Nishimura et al. 2009; Yesbolatova et al. 2020). Phosphorylation-dependent protein degradation can couple cellular signaling directly to changes in protein abundance. This has motivated engineering of a tyrosine-phosphorylation-based synthetic signaling network in human cells, as well as serine/threonine-phosphorylation-based phospho-regulons responding to MAPK Fus3 activated by mating signaling in yeast (Gordley et al. 2016; Yang et al. 2025). While these engineered networks rely on either synthetic kinases or use inductors that arrest proliferation, developing modular and predictable systems that exploit endogenous signaling pathways remains challenging (Kõivomägi, Valk, Venta, Iofik, Lepiku, Morgan, et al. 2011; Nash et al. 2001).

Cyclin-dependent kinases (CDKs) offer a promising framework for engineering such systems because their activity changes dynamically throughout the cell cycle to drive controlled temporally resolved phosphorylation of hundreds of proteins (Loog and Morgan 2005; Swaffer et al. 2016; Pirincci Ercan et al. 2021). CDK substrate recognition is influenced by phosphorylation sites, cyclin docking motifs, and interactions with the phospho-adaptor protein Cks1. These features affect phosphorylation rate and degradation timing in the cell cycle, allowing substrates to respond at different kinase activity thresholds (Örd, Möll, et al. 2019). Phosphorylation by Cdk1 can affect cell cycle proteins in multiple ways, including controlling substrate protein stability, localization, and protein-protein interactions (Nash et al. 2001; Costanzo et al. 2004; Tanaka et al. 2007).

The intrinsically disordered N-terminus of the yeast protein Far1 contains CDK-responsive phospho-degron elements and associated docking motifs that influence its phosphorylation and degradation (Henchoz et al. 1997; Bhaduri and Pryciak 2011). Previous work has shown that modifying these motifs can alter Far1 degradation timing in the cell cycle, providing a basis for a modular protein stability regulator (Faustova et al. 2021). Similarly, the timing of Sic1 phosphorylation-controlled degradation was shown to be encoded by the pattern of linear motifs, while Cdk1-phosphorylation-dependent localization signals in Dna2 regulate its nuclear accumulation during the cell cycle, with the extent of nuclear accumulation affected by the optimality of motifs mediating interaction with the CDK complex (Nash et al. 2001; Kosugi et al. 2009; Kõivomägi, Valk, Venta, Iofik, Lepiku, Morgan, et al. 2011; Faustova et al. 2022). Combining these modules may enable regulatory systems that integrate protein localization, degradation, and kinase-activity thresholds.

CDK activity and cyclin abundance vary with cell-cycle progression and in response to physiological signals (Wittenberg et al. 1990; Valk and Loog 2013). During yeast cultivation, increasing culture density and glucose limitation is associated with changes in cell-cycle distribution and growth state (Saldanha et al. 2004; Miles et al. 2013). Because Far1 degron phosphorylation and degradation dynamics depend on cyclin-CDK docking, these culture-dependent changes may affect CDK-responsive degron performance (Faustova et al. 2021). Understanding this relationship is important for developing robust regulatory systems and assessing their potential biotechnological applications.

Here, we investigate CDK-responsive phospho-degrons as modular tools for controlling protein stability and gene expression in yeast. We modify Far1-derived degrons to generate distinct degradation profiles and assess their ability to regulate the transcriptional regulators VP-EL222 and TetR. We also combine Dna2-derived phosphorylation-dependent localization modules with cyclin E- and Sic1-based degrons to investigate sequential nuclear import and degradation and to encode distinct CDK-activity thresholds within a single protein. Finally, we examine how culture density and cell-cycle distribution affect degron performance. Together, these experiments establish a framework for engineering post-translational regulatory circuits that respond to endogenous CDK activity and cellular physiology.

## Results

### Phospho-degron tags enable predictable CDK-dependent control of protein stability

The N-terminal intrinsically disordered region (residues 1–150) of Far1 contains a CDK-regulated SCF^Cdc4^ diphosphodegron flanked by Cks1 and cyclin docking motifs that influence the degron’s phosphorylation dynamics (Faustova et al. 2021). To test whether Far1 degron modules can regulate the abundance of different proteins, we used two transcriptional regulators: the light-activated transcription factor VP-EL222 (Benzinger and Khammash 2018) and the tetracycline-dependent repressor TetR (Dingermann et al. 1992).

For this, we fused four modified Far1 degron variants to VP-EL222. The variants differed in their cyclin docking motifs and Cks1 priming sites (**Fig. 1A**), aiming to obtain degrons that would be phosphorylated at different stages in the cell cycle (Faustova et al. 2021). We monitored YFP expression driven by VP-EL222 using time-lapse microscopy to provide an indirect readout of the abundance of the Far1-tagged activator (**Fig. 1B**).

**Figure 1.**
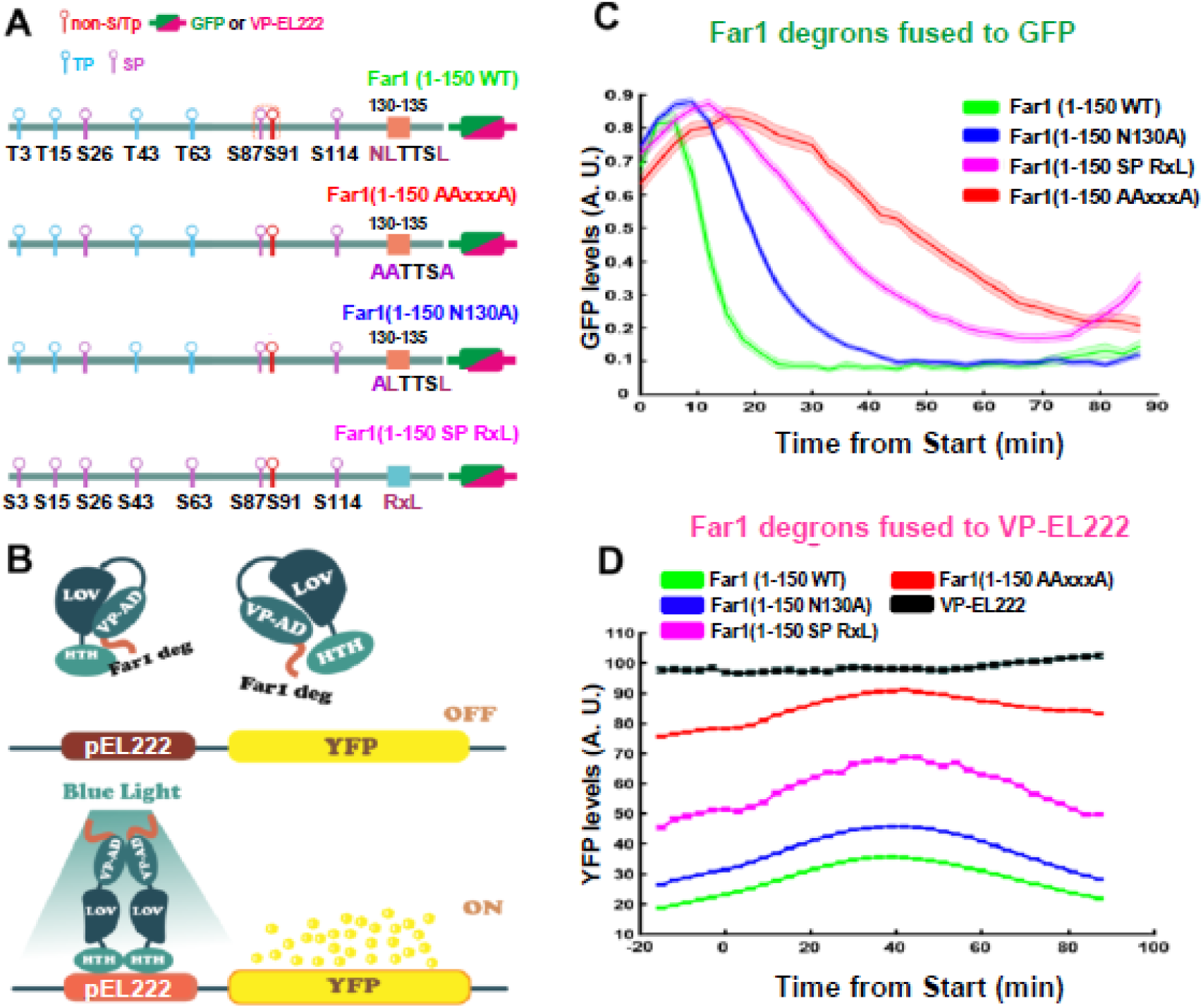
Characterization of phospho-degron tags for controlled VP-EL222 expression and output. (**A**) Schemes of Far1(1-150)-based degron tags used for fusion with GFP and VP-EL222. (**B**) Overview of the VP-EL222 light-activated transcription factor used to drive YFP expression controlled by the pEL222 promoter. Following blue-light activation, the VP-EL222 transcription factor binds the pEL222 promoter and induces YFP expression. Fusion of Far1-based degrons to the transcription factor mediate its phosphorylation and subsequent degradation. The degradation rate depends on the phospho-degron type and the efficiency of phosphorylation and protein turnover, thereby modulating YFP expression. (**C**) Degradation profiles of the indicated Far1(1-150)-GFP variants in yeast cell cycle measured in time-lapse fluorescence microscopy. Data is from Faustova et al., 2021. Start was determined by nuclear export of 50% Whi5-mCherry. (**D**) YFP expression profiles over the cell cycle when expressed from pEL222 promoter, activated by VP-EL222 transcription factor fused to different Far1-based phospho-degron variants.

Previous microscopy experiments showed that wild-type Far1(1–150) is rapidly degraded at cell-cycle onset, reaching half of its nuclear level in approximately 11 min and near-complete depletion in 20 min. By contrast, degradation of the Far1 AAxxxA mutant (with mutated Clb5 docking motif NLxxxL) was almost five times slower (**Fig. 1C**), highlighting the contribution of the NLxxxL motif to Far1 phosphorylation. The N130A mutation partially disrupts NLxxxL-mediated docking and slows degradation relative to wild-type Far1, increasing the tagged TetR half-life in the cell cycle to approximately 20 min. The RxL SP mutant, in which NLxxxL was replaced by an RxL motif and TP sites were mutated to SP to disrupt Cks1 docking, was more stable than N130A, with a 50% degradation time of approximately 35 min. Together, these variants create a graded series of cell-cycle-dependent degradation rates that can be used to tune cell-cycle-dependent protein stability.

We next tested whether these differences in degron kinetics could regulate YFP expression in cells. VP-EL222 fused to the rapidly degrading Far1(1–150) variant induced lower YFP expression than the more stable Far1(1–150 AAxxxA) variant (**Fig. 1D**). This result was consistent with their respective degradation profiles in which Far1(1–150) degraded almost five times faster than the AAxxxA mutant. Similarly, the RxL SP mutant, which degraded nearly twice as slowly as N130A, was associated with approximately twofold higher cellular YFP concentration. VP-EL222 without a Far1 degron remained stable throughout the cell cycle and consistently induced high YFP expression (**Fig. 1D**). Thus, the degradation kinetics of Far1-derived tags were reflected in the expression level of the tagged transcriptional activator. These results support the use of Far1 degrons to tune protein abundance and, consequently, reporter expression during the cell cycle.

To simplify the phospho-tags, we also investigated whether a shorter Far1 degron could retain this regulatory function. We found that deletion of the N-terminal Cks1-priming sites in Far1(85-150) caused a slight delay in degradation, which could be largely rescued by improving cyclin docking-mediated phosphorylation by decreasing the distance between the phospho-degron and the NLxxxL motif (**Fig. S1A-B**). This result suggests that degron kinetics can be further tuned by altering the spacing between motifs and phosphorylation sites and that the phospho-degron module can be encoded in less than 50 residues.

### Cdk1-controlled phosphorylation of Far1 modules regulates TetR abundance and output

To assess whether Far1 degrons can regulate a transcriptional repressor as well as an activator, we fused Far1 degron variants to TetR. This strategy was intended to control TetR stability and thereby modulate its repression activity (**Fig. 2A-B**).

**Figure 2.**
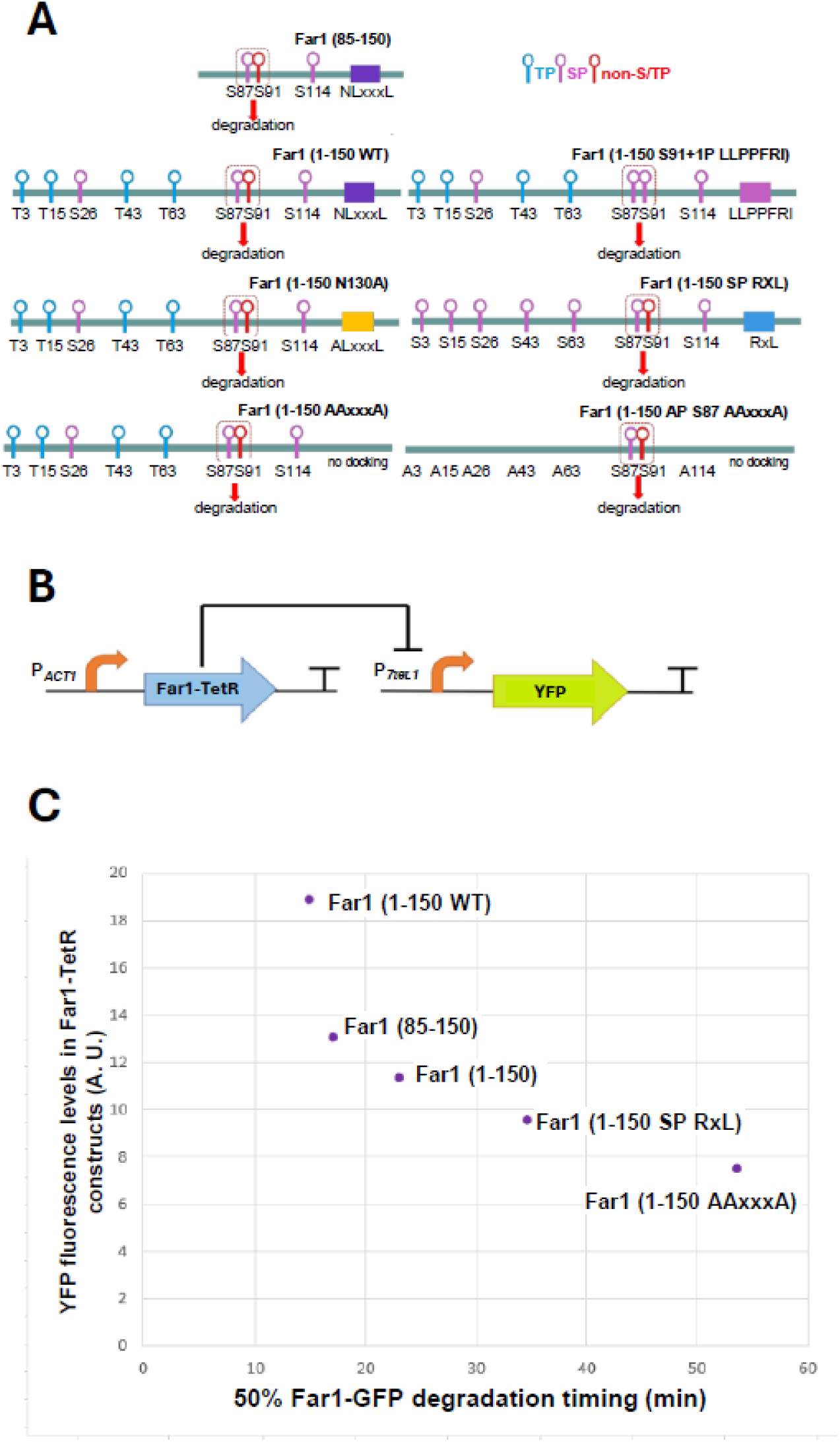
Regulating TetR transcriptional repressor output with phospho-degron tags. (**A**) Schemes of Far1(1-150)-based degron tags fused to the N terminus on TetR. (**B**) Constitutively active TetR transcriptional repressor tagged with phospho-degron inhibits expression of YFP. YFP fluorescence is negatively correlated with TetR levels, which are modulated by fusion to Far1-based phospho-degron variants. (**C**) YFP expression levels in cells expressing TetR repressors fused to the indicated degron tags (on the y-axis) and the degradation timing of these degron tags when fused to GFP in an unperturbed cell cycle (x-axis).

The degron-tagged TetR variants drove different levels of repression from the tetracycline-inducible TetO promoter (**Fig. 2C, S2**). TetR fused to the rapidly degrading Far1(1–150) degron led to high YFP expression, whereas TetR fused to the more stable Far1 AAxxxA variant showed stronger repression of YFP. The N130A and RxL SP variants produced intermediate YFP signals, consistent with their degradation profiles. These findings indicate that Far1 degron variants can tune TetR abundance and its associated promoter-repression activity.

Together, the VP-EL222 and TetR experiments show that Far1-derived degrons can regulate both transcriptional activation and repression. Their distinct degradation kinetics were reflected in the abundance and activity of two different regulatory proteins, supporting the use of these modules as adaptable components for predictable control of gene expression.

### A two-step mechanism enables degradation of cytoplasmic proteins

The phospho-degrons described above are recognized by SCF^Cdc4^ in the nucleus. Nuclear degradation through SCF^Cdc4^ is characterized in more detail than cytoplasmic degradation mediated by SCF^Grr1^ (Quilis and Igual 2017). As many metabolic enzymes used in microbial cell factories are cytoplasmic, we sought to couple phospho-degron-mediated protein degradation with regulated nuclear import. For this purpose, we used the bipartite nuclear-localization module from Dna2 (Faustova et al. 2022; Kosugi et al. 2009), which translocates from the cytoplasm to the nucleus, where the tagged protein can be targeted for SCF^Cdc4^-dependent degradation.

We tested a modified mammalian cyclin E phospho-degron (Ye et al. 2004) (LLTPPRSP) at either the N- or C-terminus of the wild-type Dna2(1–100) module, using an inactive degron (LLAPPRAP) as a control (**Fig. 3A**). The construct with the C-terminal degron showed the lowest nuclear concentration, suggesting rapid degradation following nuclear import (**Fig. 3B**). The protein with the N-terminal degron also degraded more rapidly than the construct with the mutated degron and constructs without the degron, although its nuclear accumulation at the onset of DNA replication remained detectable. Measurements of total cellular GFP showed substantially lower abundance for the C-terminal degron construct and a modest reduction for the N-terminal construct relative to the controls (**Fig. 3C**).

**Figure 3.**
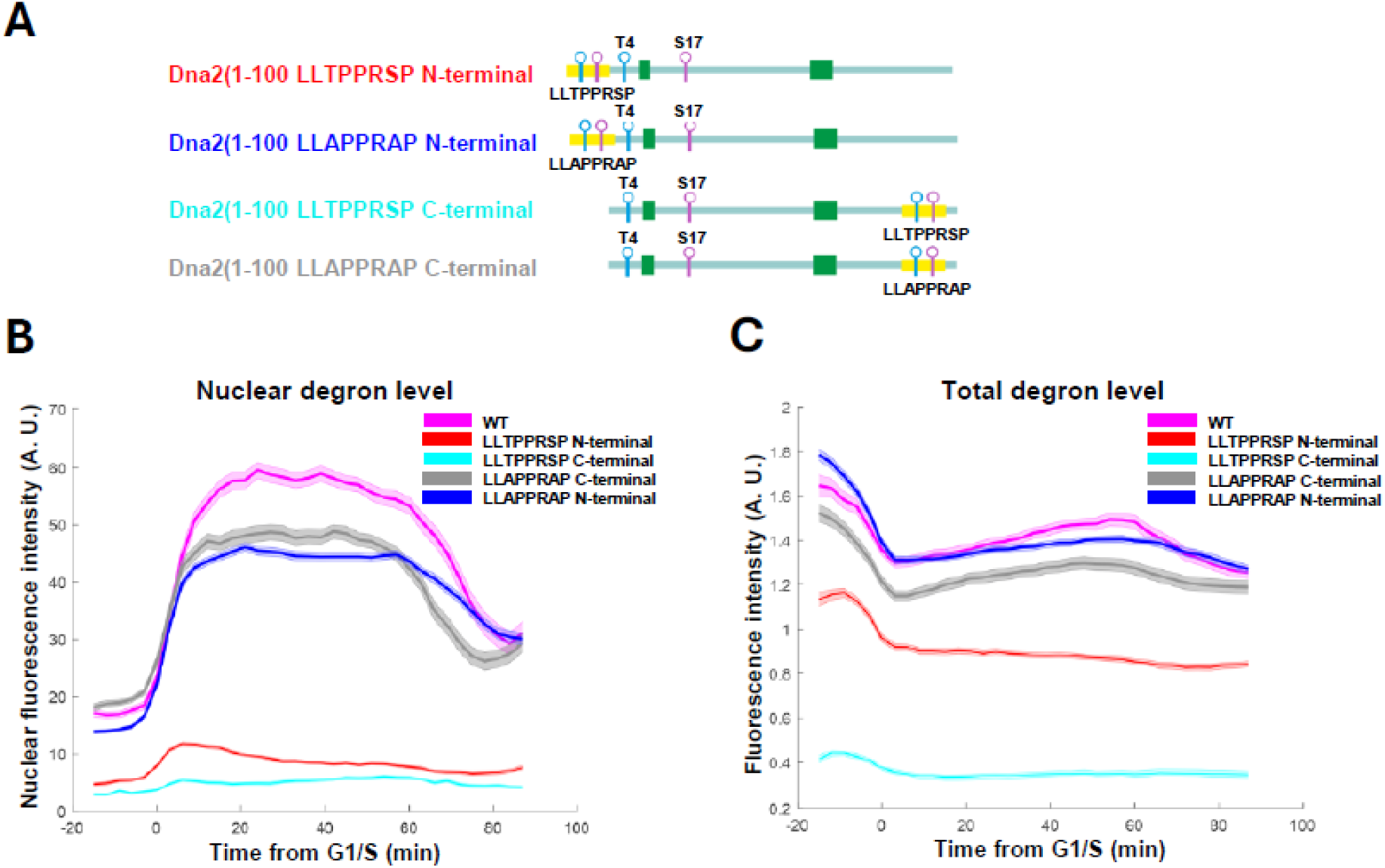
Combined phosphorylation-controlled nuclear import and degradation. (**A**) Schemes of Dna2(1-100)-based nuclear shuttling tags. Following nuclear import, the phosphorylation of LLTPPRSP sequence (a human cyclin E-derived SCF^Cdc4^ phospho-degron) promotes protein turnover through SCF^Cdc4^-mediated ubiquitylation and proteasomal degradation. (**B**) Nuclear Dna2(1-100)-EGFP fluorescence levels measured in time-lapse microscopy using the Mcm2/3-mCherry sensor to synchronize individual cells at the G1/S transition. Dna2-based nuclear shuttling tags were fused to EGFP at either the N or C terminus, thereby affecting its localization and degradation rate. (**C**) Total levels of Dna2(1-100)-EGFP variants throughout the cell cycle in time-lapse fluorescence microscopy. The data shown is average fluorescence levels from a population of cells synchronized at G1/S transition.

These results support a two-step strategy for regulating cytoplasmic proteins: phosphorylation of the Dna2 module promotes nuclear import, after which the cyclin E-derived degron mediates rapid SCF^Cdc4^-dependent degradation. The observed differences between the N- and C-terminal configurations further indicate that degron placement can influence the outcome of this combined localization-and-degradation design.

### Multiple CDK phosphorylation outputs with different kinase activity thresholds can be encoded in one protein

Although phosphorylation sites in natural CDK substrates are often phosphorylated at similar times during the cell cycle, giving a substrate a single effective CDK-activity threshold (Swaffer et al. 2016), we asked whether a single protein could encode multiple Cdk1 phosphorylation outputs with distinct timing. Cyclin E-derived degrons provide rapid degradation, whereas engineered Sic1 degrons exhibit a range of phosphorylation thresholds (Örd, Venta, et al. 2019). We reasoned that combining Dna2 localization modules with Sic1 degrons activated at higher CDK thresholds could permit transient nuclear accumulation before degradation.

To test this possibility, we fused two Sic1 variants to Dna2(1–100) (**Fig. 4A**). The first contained Sic1(1–130) with two RxL motifs to facilitate S phase cyclin binding. The second also contained four additional amino acids downstream of the Cks1 priming site T33 (T33+4), a modification designed to reduce the phosphorylation rate of the phospho-degron (Örd, Venta, et al. 2019).

**Figure 4.**
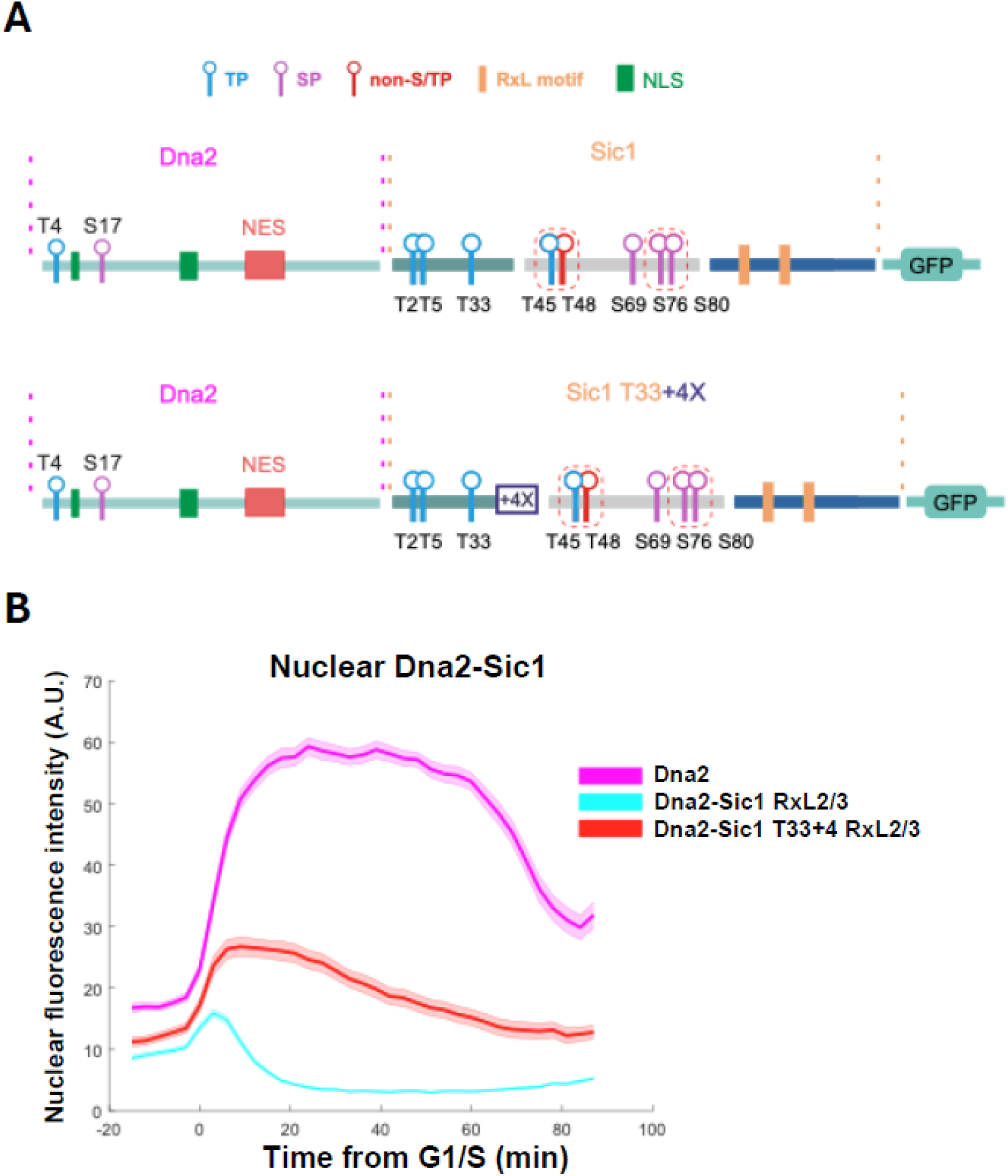
Encoding phospho-tags with different CDK activity thresholds for nuclear import and degradation. (**A**) Schemes of Dna2(1-100)-based nuclear shuttling tags fused to SCF^Cdc4^ degron sequences from Sic1. In the T33+4 degron variant, four amino acids were added downstream to the Cks1 priming site (T33+4) to reduce the phosphorylation rate of the phospho-degron (Örd, Möll, et al. 2019). (**B**) Nuclear fluorescence levels of Dna2-Sic1-EGFP modules measured by time-lapse microscopy. Average fluorescence intensity of cells synchronized at the G1/S transition using an Mcm2/3-mCherry sensor is shown.

Both Dna2-Sic1 constructs degraded more slowly than the construct with the cyclin E degron described above (**Fig. 4B**; compare with **Fig. 3B**). Their nuclear accumulation was evident shortly after the G1/S transition (**Fig. 4B**). The first Sic1 variant showed less nuclear accumulation and faster degradation than the T33+4 variant, which was degraded more slowly. This difference is consistent with the expected effect of delayed phospho-degron phosphorylation in the construct with increased spacing between T33 and the degron (T33+4). At G1/S, the Dna2-Sic1 fusion constructs began to decline, whereas wild-type Dna2 showed an opposing pattern (**Fig. 4B**).

These results indicate that Sic1 degron architecture can alter the timing of degradation and that different CDK activity thresholds can be incorporated into a single protein. The combined localization and degron modules therefore provide a means of engineering multiple temporally distinct responses to CDK activity.

### Culture density affects the stability of CDK-responsive degron tags

Previous studies on phosphorylation of Cdk1 substrates have generally examined cells during logarithmic growth at optical densities (OD) below 1. As microbial cell factories, where the designed phospho-modules could be used, are often cultivated to higher densities, we investigated whether degron regulation changes with culture OD. We also examined cell-cycle distribution and the abundance of Clb5, a major cyclin involved in Far1 degradation (Faustova et al. 2021).

We grew yeast expressing endogenous Clb5 tagged with 3HA in YPD to different ODs and collected samples for flow-cytometric analysis of cellular DNA content and for western blotting to measure Clb5 abundance. As expected, SYBR Green staining showed a progressive increase in cell counts with increasing culture OD (**Fig. S3A**). Below OD 1, most cells were in G2/M phase, and by OD 4, the G1 fraction had increased modestly. The cultures reached a maximum OD of approximately 16, at which point proliferation ceased, likely due to depletion of glucose in the medium.

Similar to the shift to the G1 phase, cyclin Clb5 abundance also decreased with increasing culture OD (**Fig. S3B**). While Clb5 abundance remained largely unchanged up to OD 6.6, it was substantially lower at OD 15. Thus, increasing culture density was associated with a shift towards G1 and a reduction in Clb5 abundance, suggesting that CDK-responsive degron activity also changes with culture state.

To examine degron stability during batch cultivation, we grew yeast strains expressing different Far1-based degron tags fused to TetR in YPD and collected samples at different ODs. TetR expressed without a Far1 degron or fused to Far1(1–150 AP S87 AAxxxA) showed no visible YFP signal throughout the experiment (**Fig. 5A-B**), consistent with high expression of the TetR repressor and the low YFP levels measured by time-lapse microscopy (**Fig. 2**). The other Far1 degron variants showed YFP expression patterns similar to those in Fig. 2 until OD 8, after which the YFP signal began to disappear across strains. Far1(1–150 SP RxL) and Far1(1–150 AAxxxA) showed no visible YFP signal at OD 8. At this density, flow cytometry also showed a lower G2-phase fraction and a higher G1-phase fraction, suggesting a decrease in CDK activity. This resulted in more stable expression of the degron-fused TetR, followed by a loss of the YFP signal (**Fig. 5A**).

**Figure 5.**
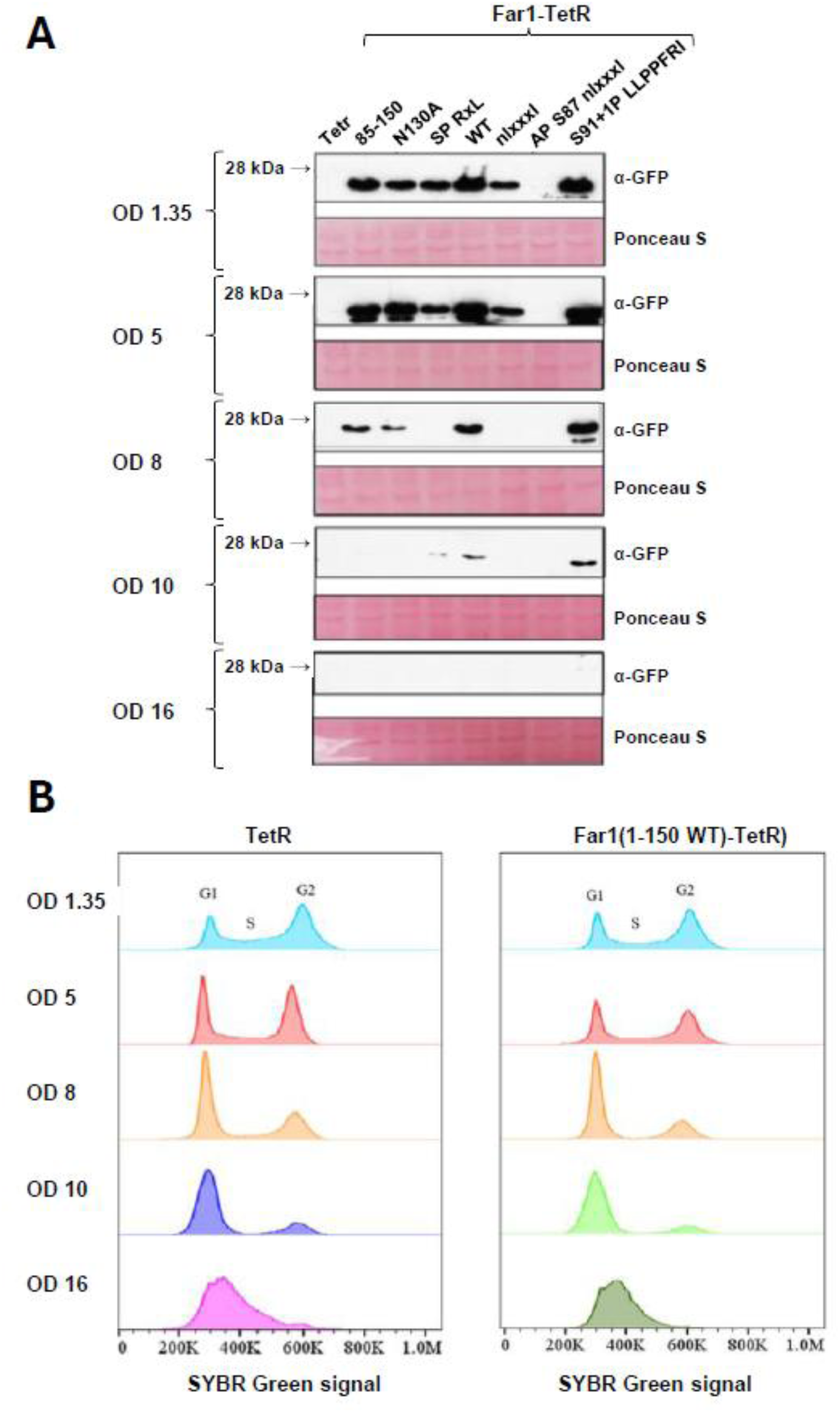
OD-dependence of phospho-tags running on endogenous cell cycle machinery. (**A**) Immunoblots showing the YFP expression controlled by degron-tagged TetR at different culture ODs. To detect YFP protein, membranes were probed with anti-GFP antibodies. Membranes stained with Ponceau S are shown as loading control. A representative experiment of two independent replicates is shown. (**B**) Flow cytometry profiles of the same cultures showing cell cycle profiles with SYBR Green DNA staining.

Overall, these measurements associate increasing culture density with changes in cell cycle distribution, reduced Clb5 abundance, and altered levels of degron-tagged proteins. The results suggest that culture state is an important variable when evaluating the performance of CDK-responsive degrons.

## Discussion

Achieving precise, dynamic control of protein abundance across time and physiological states remains a central challenge in cell biology and synthetic biology (R. P. Chen et al. 2019; Natsume and Kanemaki 2017). Although transcriptional regulation has been extensively engineered into predictable genetic circuits (Nielsen et al. 2016; Elowitz and Leibler 2000), the development of composable post-translational control systems has only recently begun. Protein-level circuits and logic have been built using engineered proteases and *de novo* designed protein modules (Gao et al. 2018; Z. Chen et al. 2020). In yeast, modular phosphorylation tags controlled by the MAP kinase Fus3 have been used to engineer synthetic kinase-substrate relationships (Gordley et al. 2016). Here, we establish CDK-responsive phospho-degrons as a modular platform for regulating protein stability and demonstrate how their properties can be used to construct dynamic regulatory behaviors.

A key finding is that Far1-derived degron variants provide tunable control over protein stability. By modifying cyclin docking motifs and Cks1-dependent priming sites, we generated variants with distinct degradation kinetics. These differences were reflected in protein abundance and reporter output. Similar behavior was observed in both a transcriptional activator and a repressor, supporting the transferability of Far1 degrons across different protein contexts.

Our results also show that CDK-responsive modules can be combined to encode different kinase activity thresholds within a protein. Pairing Dna2-derived localization modules with degrons of varying sensitivity produced architectures in which nuclear import and degradation occurred sequentially (Kosugi et al. 2009; Faustova et al. 2022). Sic1-based degrons supported transient nuclear accumulation before degradation, whereas the cyclin E-derived degron promoted more rapid turnover (Ye et al. 2004). These observations are consistent with multisite phosphorylation models in which substrate responses depend on the number, affinity, and spatial arrangement of phosphorylation sites (Nash et al. 2001; Kõivomägi, Valk, Venta, Iofik, Lepiku, Balog, et al. 2011; Kõivomägi et al. 2013; Örd, Möll, et al. 2019). They also demonstrate how these principles can be applied to engineer responses to endogenous kinase activity.

The performance of CDK-responsive degrons was also associated with cellular physiology. As culture density increased, cells accumulated in G1 and the abundance of cyclins declined. These changes coincided with altered levels of degron-tagged proteins, consistent with reduced CDK-dependent phosphorylation and degradation at higher culture densities. This coupling to growth state could provide a route to autonomous regulation during batch cultivation. However, our results show only an association between culture density, cell-cycle state, and degron behavior, and the contribution of each factor will require further investigation.

This growth-state dependence may have implications for biotechnology and metabolic engineering. Microbial cell factories often face a trade-off between biomass accumulation and production of target compounds (Venayak et al. 2015; Lalwani et al. 2018), because expressing heterologous pathways during growth can impose a metabolic burden (Wu et al. 2016). CDK-responsive degrons could potentially support growth-phase-dependent control of enzyme abundance: degradation during active growth might limit pathway burden, whereas reduced degradation as cells enter a later growth state could permit enzyme accumulation (**Fig. 6**). Inducible protein degradation and cell-cycle control have already been used in yeast to redirect flux and decouple growth from production (Lu et al. 2021; Bezold et al. 2023). Such strategy may be useful for pathways in which flux must be balanced between growth and production, including the mevalonate pathway (Pitera et al. 2007; Peng et al. 2017). These applications remain prospective and will require direct testing in production strains and pathways.

**Figure 6.**
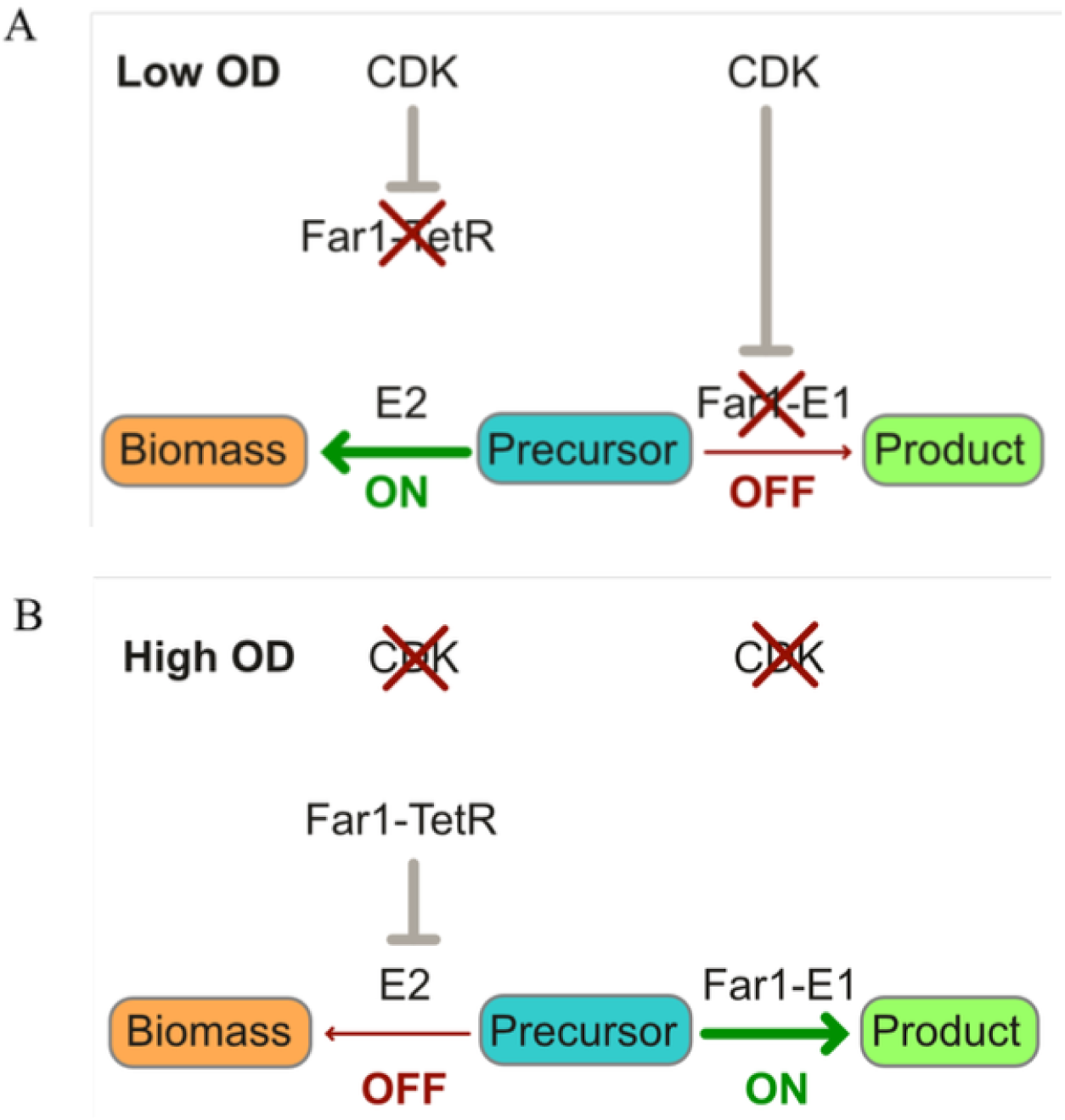
Far1-based phospho-degron tags can be used as both ON and OFF switches to regulate the levels of desired proteins E1 and E2. (**A**) The phospho-degron system activity at low OD: Cdk1 is active; hence, Far1-TetR is degraded, and E2 (an essential metabolic enzyme allowing proliferation) is expressed, leading to biomass accumulation. Simultaneously, E1 tagged with a phospho-degron module is degraded, and product synthesis is low. (**B**) System behavior at high OD: Far1-based phospho-degron tags are stabilized due to the absence of Clb5; hence, Far1-tagged E1 protein is expressed, increasing the amount of the final product. In parallel, Far1-TetR is also stabilized, allowing accumulation of TetR that regulates E2. Thus, the expression of E2, and as a result, biomass accumulation is suppressed.

The modularity of phospho-degron tags may also enable combinatorial regulation of multiple proteins in a cell. Tagging enzymes or regulators with degrons of different strengths could provide a means of tuning pathway flux without changing promoter architecture (Peng et al. 2017, 2018). Combined with genome engineering approaches such as CRISPR-based tagging (DiCarlo et al. 2013), this strategy could support coordinated protein-level regulation of metabolic pathways. Engineered cyclin-CDK modules or inducible kinase systems may further expand the dynamic range and programmability of such circuits.

In summary, this study establishes CDK-responsive phospho-degrons as flexible tools for controlling protein stability and engineering dynamic regulatory behaviors. By exploiting multisite phosphorylation, these modules can tune protein abundance, support sequential localization and degradation, and couple protein regulation to cell-cycle state and culture conditions. They provide a foundation for further development of synthetic circuits and for exploring post-translational control in biotechnology.

## Acknowledgements

This work was funded by the Estonian Research Council grant PRG2149. This work received co-funding from the European Union’s Horizon Europe research and innovation programme under Grant Agreement No. 101060066 (DIGIBIO). It was also supported by the European Union through the European Regional Development Fund (Center of Excellence for Bioengineering, project No. 2021-2027.1.01.24-0451) and by the Estonian Ministry of Education and Research. Views and opinions expressed are, however, those of the author(s) only and do not necessarily reflect those of the European Union or the European Research Executive Agency (REA). Neither the European Union nor the granting authority can be held responsible for them. The authors would like to thank Regina Virre for her excellent technical assistance and dedicated support in maintaining the laboratory infrastructure throughout this study. The authors would like to thank Mart Loog for making this project possible.

## Author contributions

Conceptualization: M.Ö., I.F.; Funding acquisition: I.F.; Investigation: M.L., A.A., L.B., S.S., M.Ö.; Methodology: M.L., A.A., L.B., S.S., E.V.; Project administration: M.Ö., I.F.; Resources: E.V.; Supervision: M.Ö., I.F.; Writing – original draft: M.Ö., N.S.; Writing – review & editing: M.L., A.A., L.B., S.S., E.V., I.F.

## Methods

### Yeast methods

*S. cerevisiae* strains were derived from W303 and are described in Table S1. Gene deletions and tagging were performed using polymerase chain reaction and homologous recombination–based approach.

### Time-lapse fluorescence microscopy

Before microscopy experiments, the cells were grown at 30°C in synthetic complete (SC) medium supplemented with 2% glucose to OD_600_ (optical density at 600 nm) of 0.2 to 0.6. The culture was pipetted onto 0.8-mm cover glass and covered with a 1-mm-thick 1.5% agarose (NuSieveTM GTGTM Agarose, Lonza) SC/glucose pad. The sample was kept under the microscope for 1 hour before starting the experiment. The experiments were carried out using Zeiss Observer Z1 microscope with a 63×/1.4 numerical aperture oil immersion objective and Axiocam 506 mono camera (Zeiss). The sample was kept at 30°C with Tempcontrol 37-2 digital (PeCon). The cells were imaged for phase contrast, GFP, and mCherry with 3 × 3 binning every 3 min. The experiments were up to 8 hours long, and up to 12 positions were tracked using ZEN software (Zeiss), an automated stage, and Definite Focus. Filter Set 61 HE (Zeiss) in combination with Colibri 470 and Colibri 540-580 light-emitting diode (LED) modules with exposure times of 15 and 450 ms was used for imaging of sfGFP and mCherry, respectively. The Colibri LED modules were used at 25% power.

Image segmentation, cell tracking, and quantification of fluorescence signals were performed using MATLAB (The MathWorks Inc.). To analyze nuclear levels of the fluorescent proteins, the brighter region of the cell was segmented as the nucleus, and the cytoplasm was treated as background signal, as the same amounts of the fluorescent proteins have very low signals when diluted into the cytoplasm. Also, as the cytoplasmic fluorescent protein signal overlaps with the nuclear signal, the average cytoplasmic signal is subtracted from the average nuclear signal to obtain the final nuclear fluorescence intensities shown on the plots. All plots with microscopy data contain data from at least two biological replicate experiments. No cells or replicate experiments were excluded from the analysis. The data are plotted as the average nuclear fluorescence intensities for a cell population across multiple experiments. The error bars are ±SEM. No sample size pre-calculations were made. Fifty cells were considered sufficient for the analysis because replicate experiments with this sample size showed very little variation.

### Western blot

The yeast cultures were grown at 30°C in yeast extract, peptone, and dextrose (YPD) medium. Cells from 5 ml of the culture were collected and flash-frozen in liquid nitrogen every 10 min. The transfer to a nitrocellulose membrane was performed using the standard semidry transfer with Pierce G2 Fast Blotter (Thermo Fisher Scientific). Purified anti-HA.11 epitope tag antibody (1:500) (clone 16B12, BioLegend, catalog no. 901501, RRID:AB_2565006) and horseradish peroxidase–conjugated anti-mouse antibody (1:7500) from LabAs, Estonia were used to detect the 6HA-tagged proteins. All Western blotting experiments were done in at least two biological replicates.

### Flow cytometry

Yeast cultures were pipetted into 80% ethanol, and cells were fixed overnight in a rotator at 4°C. Fixed cells were treated with RNase A (Sigma) (1 mg/ml RNase A for 40 min at 37°C) and Proteinase K (Thermo Scientific) (200 μg/ml Proteinase K for 1h at 37°C). DNA was stained with SYBR Green I (Sigma-Aldrich) for 1 h.

