## Supplementary Figures for "CDK phospho-degron tags for dynamic protein regulation"

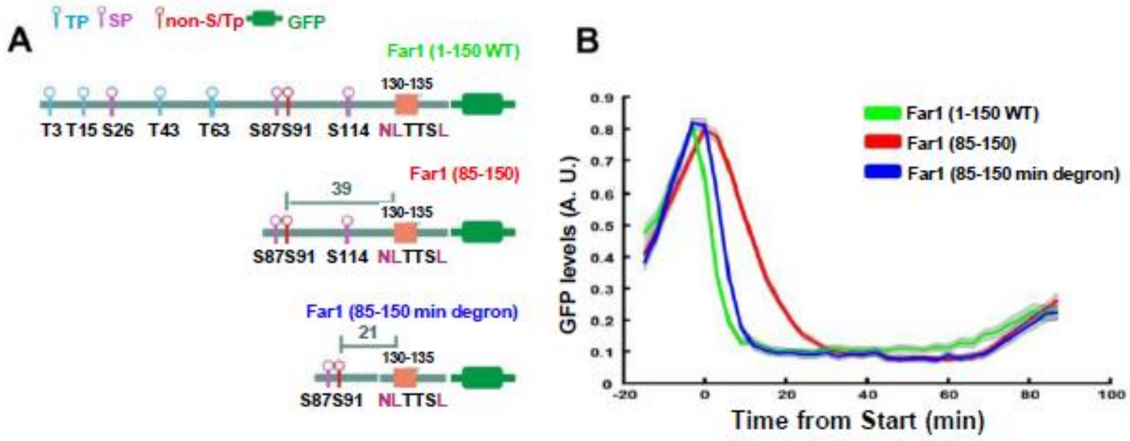


**Supplementary Figure 1. Designing a minimal variant of Far1 phospho-degron.** (**A**) Schemes of Far1-based degron tags. (**B**) Tracking of different Far1 degron-GFP protein expression profiles in the cell cycle using time-lapse fluorescence microscopy.

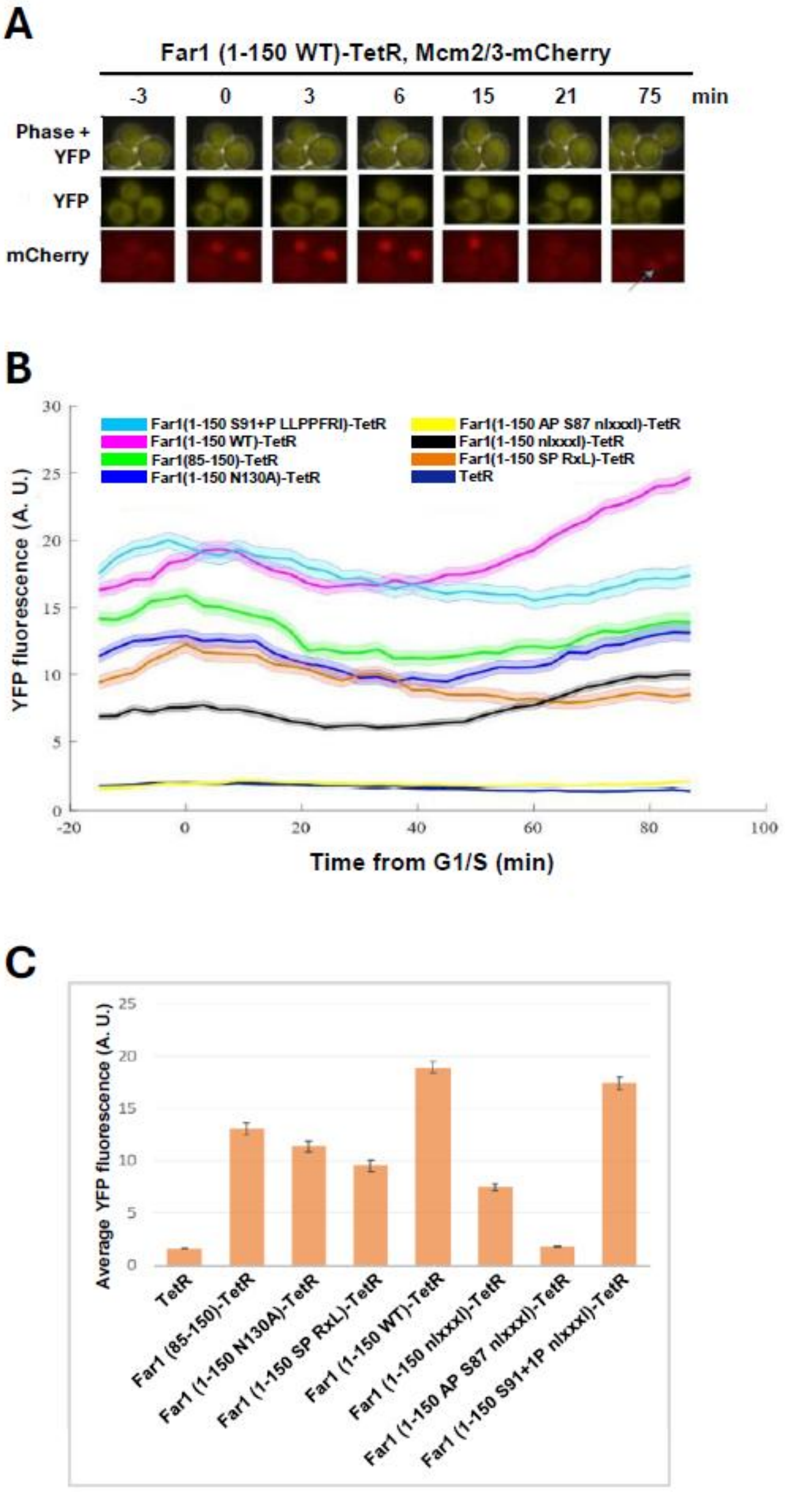


**Supplementary Figure 2. Regulation of TetR by degron tags.** (**A**) Examples of time-lapse microscopy images showing nuclear export of Mcm2/3-based cell cycle reporter (mCherry) and YFP controlled by TetR with WT degron tag. (**B**) Tracking of different YFP expression levels throughout the cell cycle in yeast cells expressing differentially tagged TetR to control the YFP expression. (**C**) Average YFP fluorescence intensity over the cell cycle. YFP expression was controlled by the indicated TetR variants.

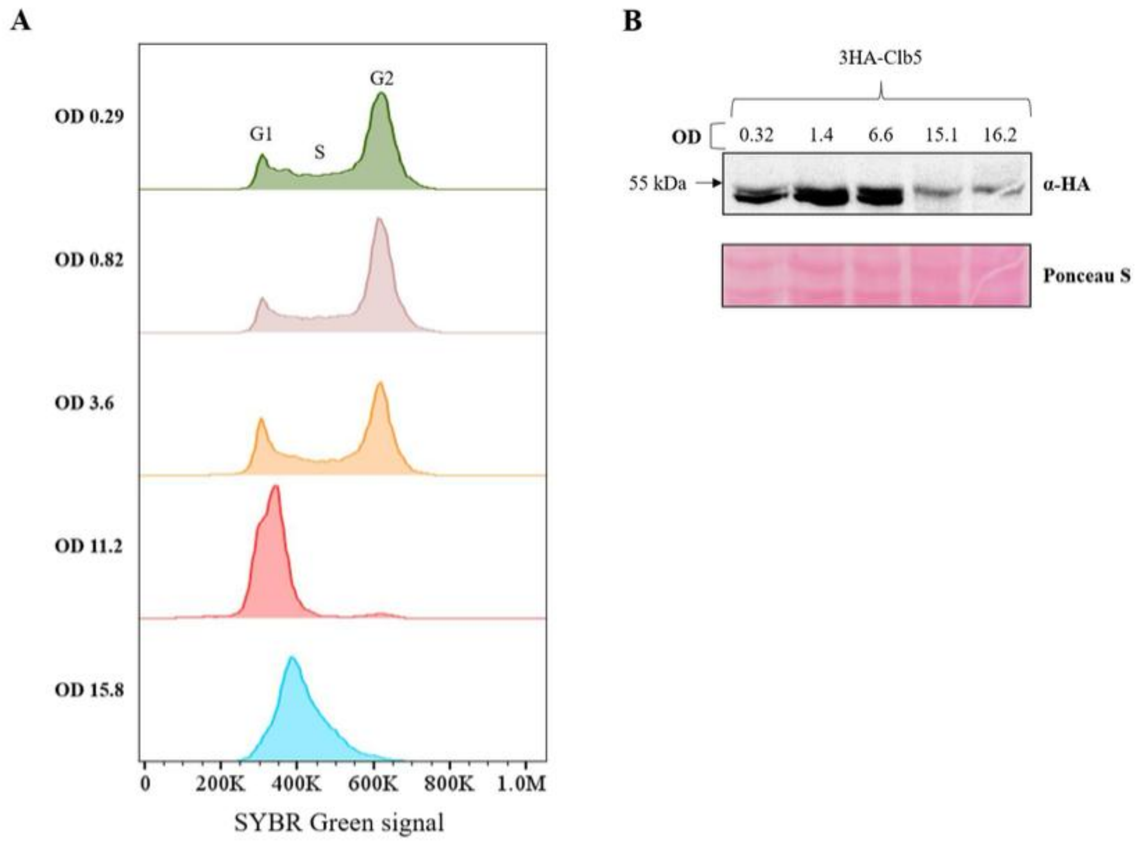


**Supplementary Figure 3. Impact of culture density on cell cycle distribution and Clb5 expression.** (**A**) Cell cycle phase distribution studied by SYBR Green staining and flow cytometry. (**B**) Western blot showing the expression of endogenous Clb5 at different culture ODs.
